# Multi-task learning from single-cell multimodal omics with Matilda

**DOI:** 10.1101/2022.06.01.494441

**Authors:** Chunlei Liu, Hao Huang, Pengyi Yang

## Abstract

Single-cell multimodal omics technologies enable multiple molecular programs to be simultaneously profiled at a global scale in individual cells, creating opportunities to study biological systems at a resolution that was previously inaccessible. However, the analysis of single-cell multimodal omics data is challenging due to the lack of methods that can integrate across multiple data modalities generated from such technologies. Here, we present Matilda, a multi-task learning method for integrative analysis of single-cell multimodal omics data. By leveraging the interrelationship among tasks, Matilda learns to perform data simulation, dimension reduction, cell type classification, and feature selection in a single unified framework. We compare Matilda with other state-of-the-art methods on datasets generated from some of the most popular single-cell multimodal omics technologies. Our results demonstrate the utility of Matilda for addressing multiple key tasks on integrative single-cell multimodal omics data analysis.

## Introduction

Recent development of single-cell multimodal omics technologies enables multiple modalities of cellular regulatory circuitry to be simultaneously profiled in individual cells ^1^. Data generated from these technologies create new opportunities for integrative analysis of cellular programs that are inaccessible from analysing each data modality alone and hence promise to provide a more holistic characterisation of cellular systems at single-cell resolution ^2^. A large number of computational methods have been developed for single-cell RNA-sequencing (scRNA-seq) data to perform tasks such as data simulation ^3^, dimension reduction ^4^ and classification of cell types ^5,6^, and feature selection ^7,8^. While methods designed for scRNA-seq data analysis can be applied to analyse RNA modality in single-cell multimodal omics data, most of them cannot take advantage of other available data modalities and therefore could not fully utilise the information embedded in such data. The lack of computational methods that can integrate across data modalities is a key issue in single-cell multimodal omics data analysis and greatly hinder biological discovery from such data ^9,10^.

Here we present Matilda, a neural network-based multi-task learning method for integrative analysis of single-cell multimodal omics data (Fig. 1a). Although previously methods developed for scRNA-seq data analysis typically address different tasks (e.g., data simulation, cell type classification) independently, a key observation in Matilda is that many common tasks in single-cell multimodal omics data analysis are closely related to each other. The modularity nature of neural networks employed in Matilda makes it well-suited for integrating multiple data modalities and performing multiple tasks in a single unified framework. For example, the data simulated by the variational autoencoder (VAE) ^11^, a key component of Matilda, can be augmented to the original data to improve cell type classification. By leveraging such relationships, Matilda simultaneously performs data simulation, dimension reduction, cell type classification, and feature selection across data modalities (Fig. 1a), therefore, achieving multiple key tasks in integrative analysis of single-cell multimodal omics data.

**Figure 1.**
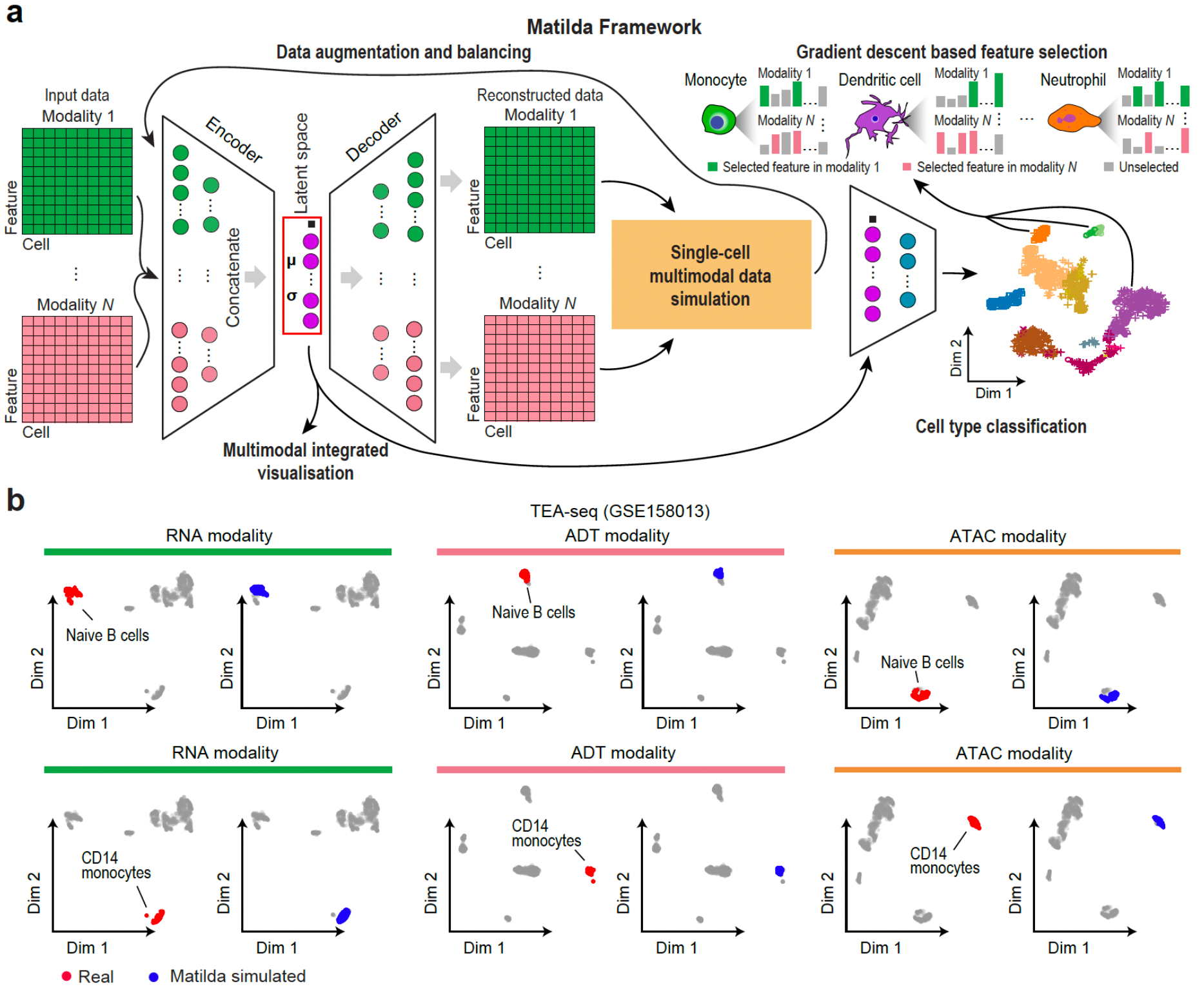
Matilda framework and multimodal single-cell data simulation. (a) Schematic summary of the main components in Matilda framework, including multimodal single-cell data simulation, data augmentation, multimodal integrated visualisation, cell type classification, and gradient descent-based feature selection. (b) UMAP visualisation of cell-type-specific simulations of RNA, ADT, and ATAC modalities in the TEA-seq dataset (GSE158013) using Matilda. Upper and lower panels show real (red) and Matilda simulated (blue) naïve B cells and CD14 monocytes, respectively.

Matilda is the first method to perform data simulation, cell type classification, and feature selection for single-cell multimodal omics data. To evaluate the performance of Matilda on multiple tasks in single-cell multimodal omics data analysis, we applied Matilda to a collection of datasets generated from popular single-cell multimodal omics technologies including those profiling three modalities using TEA-seq (gene expression [RNA], cell surface proteins [ADT], and chromatin accessibility [ATAC]) ^12^, and those profiling two modalities using CITE-seq (RNA and ADT) ^13–15^ and SHARE-seq (RNA and ATAC) ^16^. While there are currently no alternative methods available for data simulation, cell type classification, and feature selection using multiple modalities in these datasets, various methods (e.g., Sparsim ^17^ for data simulation, scClassify ^5^ for cell type classification, MAST ^8^ for feature selection) have been developed for single-cell RNA-sequencing (scRNA-seq) data and therefore can be applied using the RNA modality in these datasets. Using a range of evaluation criteria, we show that Matilda outperforms other state-of-the-art methodologies designed for various tasks using single or multiple data modalities. Our results demonstrate the utility of Matilda as the first comprehensive method for addressing multiple key tasks in single-cell multimodal omics data analysis.

## Results

### Single-cell multimodal data simulation

We applied Matilda to five recent single-cell multimodal omics datasets including a TEA-seq dataset that profiles RNA, ADT and ATAC modalities in human PBMC samples; three CITE-seq datasets that profile RNA and ADT modalities in human PBMC samples; and a SHARE-seq dataset that profiles RNA and ATAC modalities in mouse skin samples (Supplementary Fig. 1). To test if Matilda is able to simulate multimodal omics data in a cell-type-specific manner, we first visualised cells using each modality on UMAPs (Fig. 1b and Supplementary Fig. 2) and highlighted cells from representative cell types using real and Matilda simulated data. We found that Matilda not only precisely simulates each data modality in a cell-type-specific manner but also denoises the outliers in the real data, (e.g., ADT modality of B cells and CD14 cells in CITE-seq data; Supplementary Fig. 2a).

To further assess the performance of Matilda on data simulation, we compared the correlation structure of highly variable genes (HVGs) by each data modality using real data and those simulated by Matilda (Fig. 2a-c and Supplementary Fig. 3). We found that data simulated by Matilda closely resemble the correlation structure of real data across all modalities. While no other methods are currently available for simulating multimodal single-cell omics data besides Matilda, various methods have been developed for simulating from scRNA-seq data ^5^. We, therefore, compared the simulation results of Matilda on RNA modality with those generated from scGAN ^18^, a simulation method based on deep generative adversarial networks, and Sparsim ^17^, one of the best performing simulation methods based on mixture modelling ^3^. Compared to these methods, the data simulated from Matilda for the RNA modality preserve the correlation structure in the real data significantly better as quantified in Fig. 2d-f. These results demonstrate the ability of Matilda on simulating multiple data modalities in a cell-type-specific manner in single-cell multimodal omics datasets.

**Figure 2.**
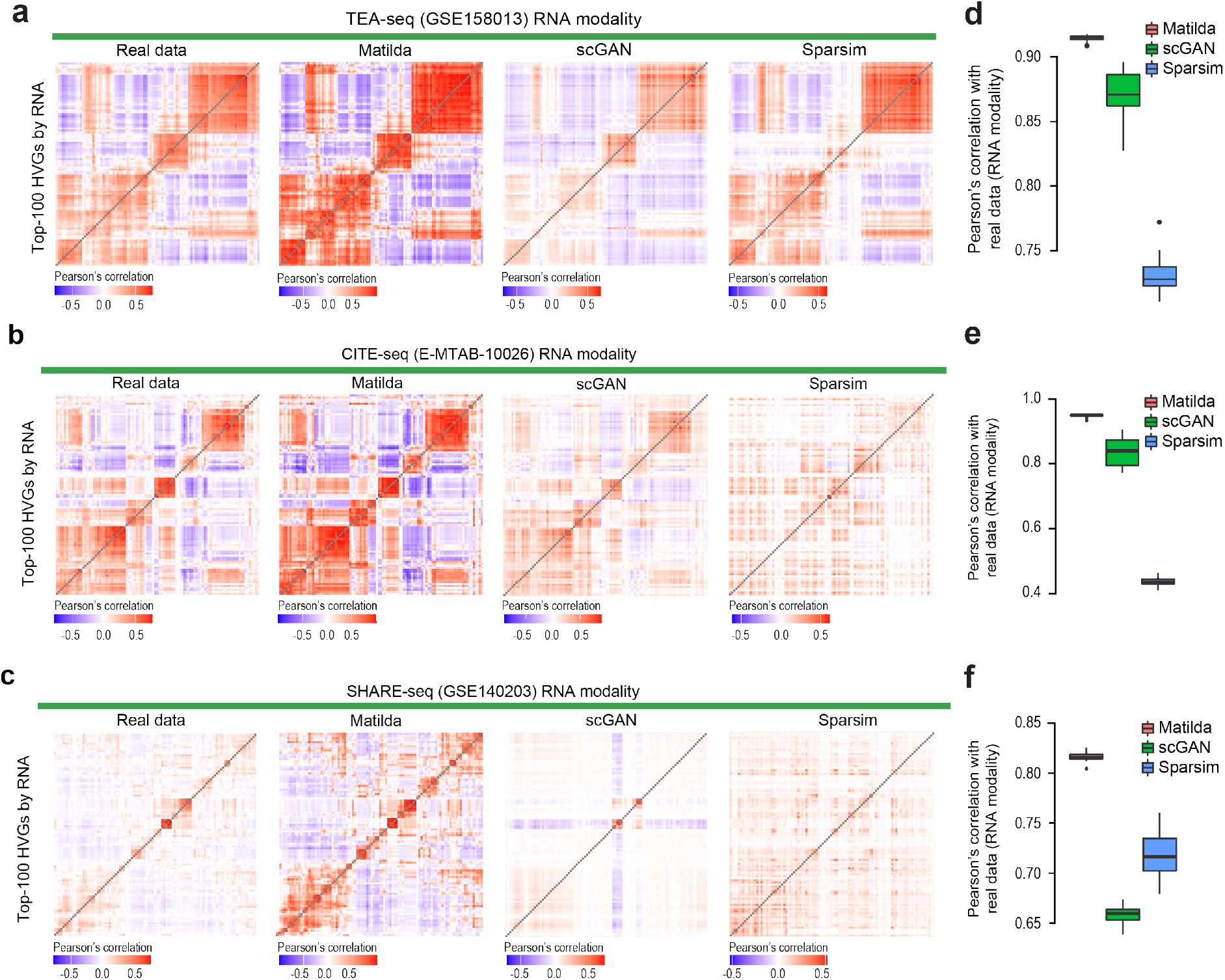
(a,b,c) Heatmap visualisation of the correlation structure of RNA modality of real and simulated TEA-seq dataset (GSE158013), CITE-seq (E-MTAB-10026) and SHARE-seq dataset (GSE140203) using Matilda, scGAN, and Sparsim. The top-100 highly variable genes (HVGs) selected from the RNA modality of the real data were used for the heatmap. (d,e,f) Pearson’s correlation of simulated data from each simulation method with real data using RNA modality for TEA-seq dataset (GSE158013), CITE-seq (E-MTAB-10026) and SHARE-seq dataset (GSE140203). Centre line, median; box limits, upper and lower quartiles; whiskers, 1.5x interquartile range; points, outliers.

### Multimodal data integration and dimension reduction

During model training, Matilda learns to combine and reduce the feature dimensions of single-cell multimodal omics data to a latent space using its VAE component in the framework (Fig. 1a). The trained VAE of Matilda thus can be used for multimodal feature integration and dimension reduction of both the training and new data. Several alternative methods are available for such tasks. These include Seurat ^14^ and totalVI ^4^, which are designed for integrating RNA and ADT modalities in CITE-seq data, and Conos ^19^ and multiVI ^20^, which are designed for integrating RNA and ATAC modalities such as these in SHARE-seq data. Comparing to these methods, we found that the dimension reduced data from Matilda shows significantly better cell type separation under UMAP projection (Fig. 3a, b).

**Figure 3.**
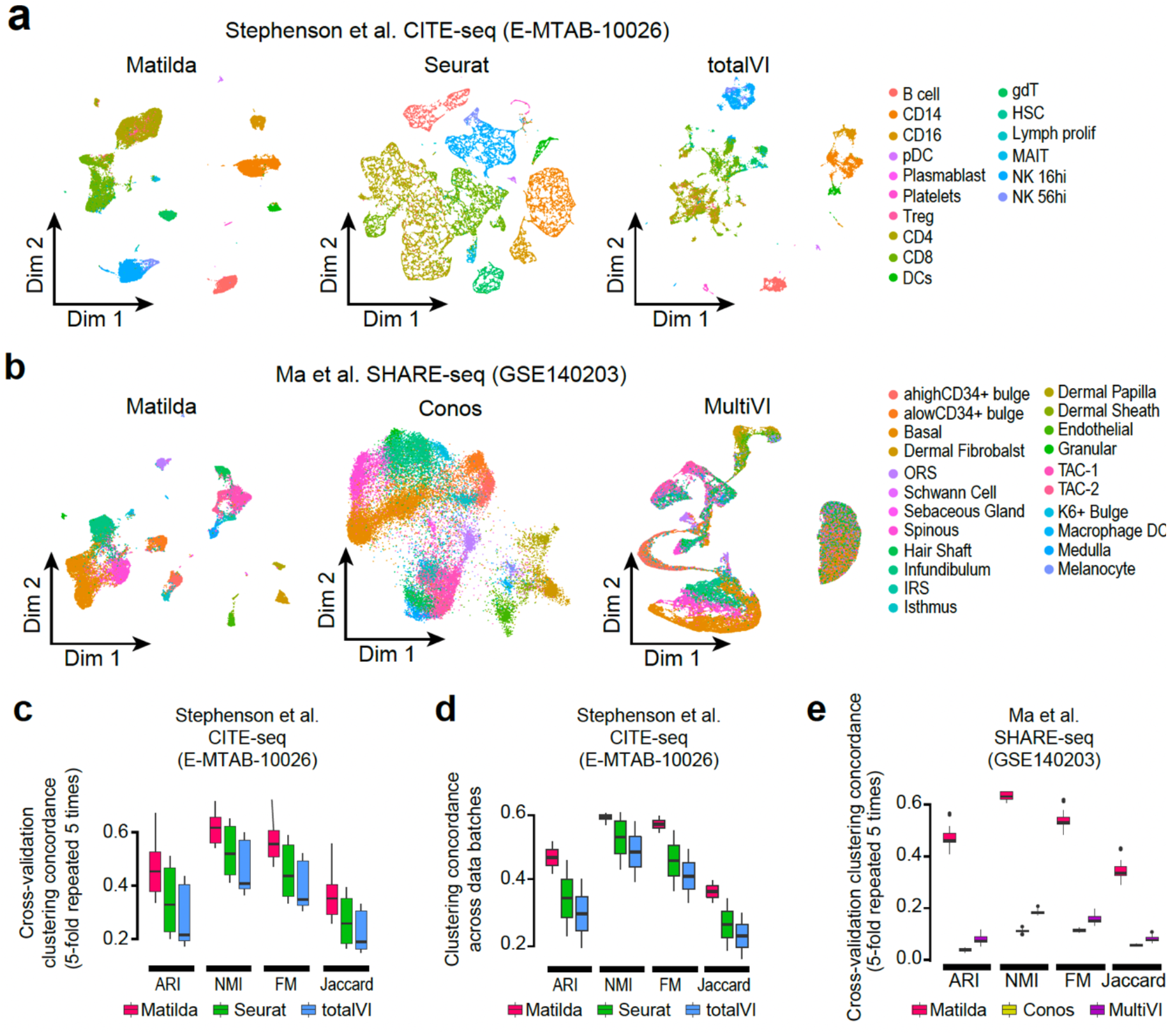
Assessment of Matilda for multimodal integrated dimension reduction and visualisation. (a, b) Visualisation and (c,d,e) quantification of joint multimodal dimension reduction results. (a) Visualisations of CITE-seq data (E-MTAB-10026) using Matilda, Seurat, and totalVI. (b) Visualisations of SHARE-seq data (GSE140203) using Matilda, Conos, and MultiVI. Cells are colour-coded by their types on the UMAPs. Quantifications were based on *k*-means clustering concordance using dimension reduced data from each method and the cell-type annotation from the original publication by ARI, NMI, FM, and Jaccard index. Either (c) 5-fold cross-validation repeated five times with different random seedings or (d) data from different batches from CITE-seq data (E-MTAB-10026) were used for capturing the variability in quantifications. (e) Quantification of *k*-means clustering concordance using dimension reduced data from Matilda, Conos, and MultiVI with cell-type annotations from the original study by ARI, NMI, FM, and Jaccard index on SHARE-seq data (GSE140203). Centre line, median; box limits, upper and lower quartiles; whiskers, 1.5x interquartile range; points, outliers.

To further quantify these visual observations, we clustered the dimension reduced data generated from each method using a simple *k*-means clustering algorithm and analysed the concordance of the clustering output with the cell type labels provided from their original studies using a panel of concordance metrics including ARI, NMI, FM, and Jaccard index (see Methods). We found that Matilda generated dimension reduced datasets led to higher clustering concordance with respect to the original cell type labels across all datasets irrespective of the metrics (Fig. 3c-e and Supplementary Fig. 4). These results demonstrate the superior performance of Matilda for integrating and reducing feature dimensions in single-cell multimodal omics data and its utility for subsequent applications such as data visualisation and clustering of cell types.

### Cell type classification using multiple data modalities

To evaluate Matilda on cell type classification using single-cell multimodal omics data, we performed both five-fold cross-validation (repeated 5 times) and training and test using different batches within each dataset (Supplementary Fig. 1b). While several methods have been developed recently for transferring cell type labels across different data modalities for single-cell multimodal omics data ^21,22^, there are currently no available methods for cell type classification by using all data modalities from such data. To this end, we resorted to comparing methods that are designed for cell type classification from scRNA-seq data by using RNA modality only ^23^. We found that Matilda classifies cells significantly more accurately across all datasets under both the cross-validation settings (Fig. 4a) and those from training and test using different batches within each dataset (Fig. 4b) than other state-of-the-art cell type classification methods that use only RNA modality. The breakdown of the classification results from training and test using each pair of data batches reveals that Matilda led to higher cell type classification accuracy across all pairs in all four datasets that contain multiple data batches (Fig. 4c).

**Figure 4.**
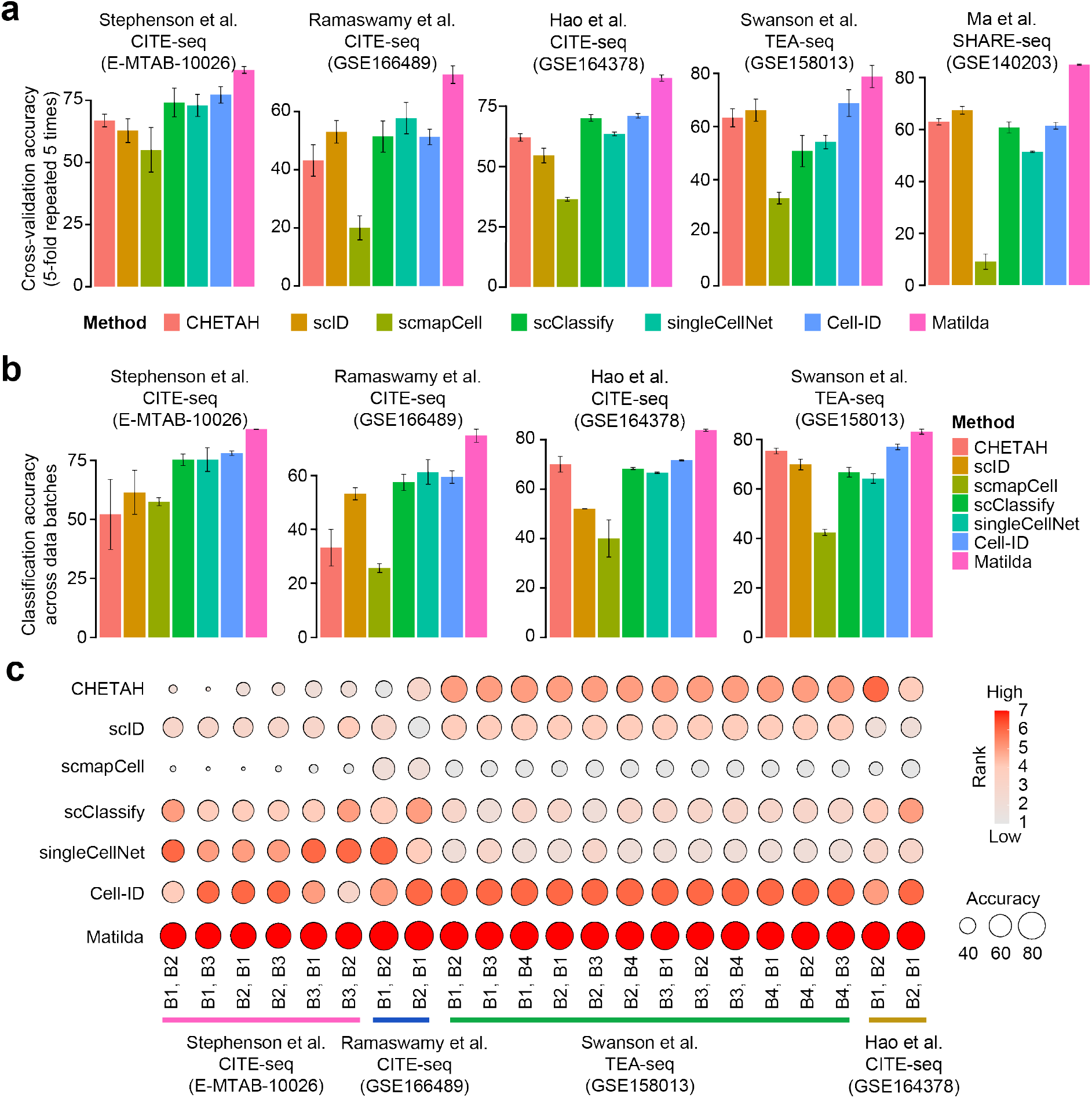
(a, b) Cell type classification of each single-cell multimodal omics data. Either (a) 5-fold crossvalidation repeated five times with different random seedings or (b) data from different batches were used for benchmarking the performance of each method. Error bar, SD. (c) Ranking summary of cell type classification accuracy across data batches for each method.

To test if the performance of Matilda is impacted by the reduced size of the training data, we performed a stratified sampling of each cell type from CITE-seq and TEA-seq datasets generated by Ramaswamy et al. ^13^ and Swanson et al. ^12^, respectively, 80%, 50%, and 20% of cells and trained each classification model using these subsampled datasets. We found that the performance of Matilda is largely maintained even when the model was trained on a small proportion of cells from the original datasets (Supplementary Fig. 5). It is worth noting that the improved cell type classification accuracy of Matilda is not a sacrifice in speed on model training or classification of test data (Supplementary Fig. 6). Since Matilda uses multi-task learning and the simulated data from the VAE component for data augmentation, we also evaluated the impact of these procedures on cell type classification accuracy. We found that, across all five datasets, multi-task learning indeed improved cell type classification than learning each task independently (Supplementary Fig. 7a), and data augmentation resulted in better performance than those without (Supplementary Fig. 7b). Together, these results demonstrate the utility of multi-task learning and data augmentation from simulation for improving cell type classification and highlight Matilda’s increased cell type classification accuracy using multimodalities compared to alternative methods that use only RNA modality.

### Feature selection from multiple data modalities

Finally, the neural network trained for cell type classification in Matilda can be used for multimodal feature selection using a gradient descent procedure ^24^ and thus can lead to the selection of cell-type-specific features across all available modalities in the datasets. Fig. 5a,b visualise top-ranked features selected by Matilda for CD14 monocytes and Naïve B cells, respectively, in each data modality in the TEA-seq dataset. The RNA and ADT expression levels and the ATAC activity of selected genes across all cell types in the dataset are shown in Fig. 5c,d. As expected, these analyses reveal that features selected by Matilda for each data modality show expression specificity towards their respective cell types, demonstrating their potential usage for characterising cell identity and their underlying molecular programs.

**Figure 5.**
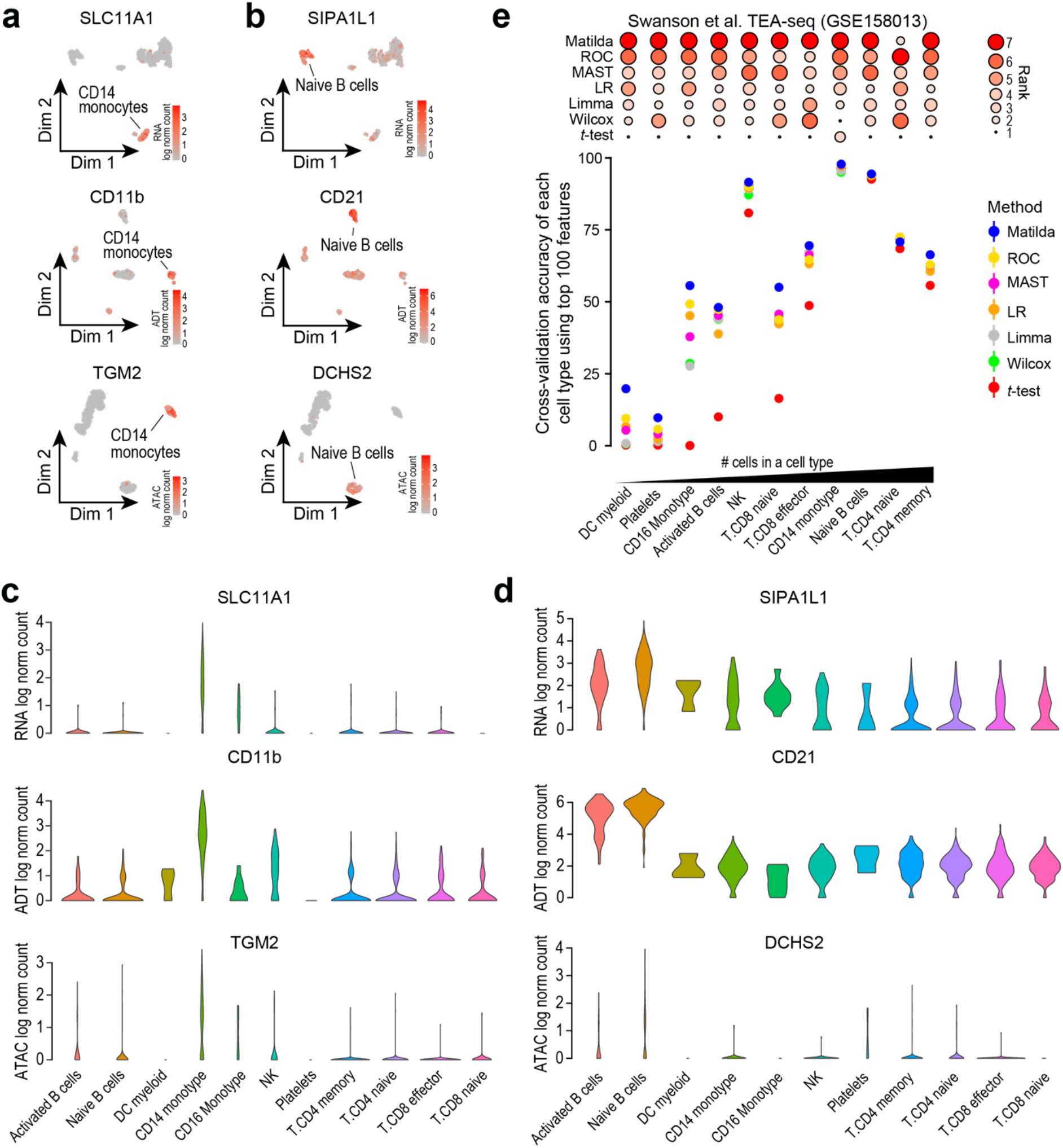
(a,b) For TEA-seq data (GSE158013), UMAPs highlight representative markers selected from each of the three modalities for CD14 monocytes and Naïve B cells. (c,d) violin plots of levels of selected markers for CD14 monocytes and Naïve B cells in their respective modalities across all cell types. (e) Classification of each cell type in TEA-seq data (GSE158013) using the cell-type-specific features selected by different methods. Cell types are arranged from low to high based on the number of cells in each cell type. Feature selection methods are also ranked based on the performance of their selected features in classifying each cell type (upper panel). Error bar, SE.

To evaluate the top cell-type-specific features selected by Matilda across multiple data modalities and those selected from RNA modality using popular methods such as t-test and limma ^7^, and those specifically designed for scRNA-seq (e.g., MAST ^8^, ROC), we compared their utility in classifying their respective cell types from all other cell types in each dataset. We found that cell-type-specific features selected by Matilda from multiple data modalities on average resulted in more accurate discrimination of their respective cell types as shown by the scatter plot and the overall rankings of methods in each dataset (Fig. 5e and Supplementary Fig. 8). Together, these results demonstrate Matilda as a useful approach for feature selection from multiple data modalities for cell type characterisation and other downstream analyses.

## Discussion

The key motivation for using multi-task learning in Matilda is that many common tasks in single-cell multimodal omics data analysis are interrelated. Learning these tasks in parallel may therefore improve the performance of the model on each individual task. Furthermore, the rationale for using neural network models in Matilda is due to their modularity which fits well with the multiple data modalities and tasks. This allows the integration of data modalities and information sharing of tasks which together enable complementary information to be extracted and hence lead to more accurate characterisations of cellular programs. With the advance in single-cell multimodal omics technologies, we expect more data modalities to become available in the near future. The modularity and flexibility of Matilda allow integration when additional modality becomes available in such data.

One common criticism of neural network-based learning models is that a large number of examples need to be provided during the training process. We demonstrated in our experiments that Matilda’s performance in cell type classification is largely maintained even with a relatively small number of cells in the training datasets. This may be due to the data simulation and augmentation component implemented in Matilda which increases the number of cells in the training datasets, especially for the rare cell types. However, dealing with cell types with an extremely small number of cells is still a challenge and may require alternative approaches.

While the current implementation of Matilda deals with datasets profiling discrete cell types, studies that look at transitional processes such as development and organogenesis create datasets with transient cell types. To analyse such datasets will require reformatting the loss function in the Matilda framework such as changing the classification component to a regression component. The potential mismatch of cell types in the training and query datasets may also have a significant impact on the performance of Matilda. A solution may be to utilise the prediction probability of the neural network for deciding whether a cell in a query dataset should be classified or not. These form the key directions for our future work.

In sum, Matilda is so far the first method for simultaneous simulation and supervised classification of cells using multiple modalities in single-cell multimodal omics data. It is also the first method for joint feature selection from multiple data modalities. Matilda addresses multiple key tasks in single-cell multimodal omics data analysis in a single unified framework.

## Methods

### Datasets and preprocessing

#### TEA-seq dataset

TEA-seq enables simultaneous single-cell profiling of transcripts, epitopes, and chromatin accessibility ^12^. The processed matrices of TEA-seq data from measuring PBMC were downloaded from the NCBI Gene Expression Omnibus (GEO) under the accession number GSE158013, with raw RNA expression, ADT expression, and peak accessibility (ATAC) measured for the same cells in four data batches. We summarised the matrix of ATAC from peak level to gene activity scores using the ‘CreateGeneActivityMatrix’ function in Seurat package ^14^. Genes with fewer than 1% quantifications across cells in each of the three modalities were removed, respectively. This resulted in a dataset with 6,310 (9,855 RNA, 46 ADT, 17,141 ATAC); 6,545 (9,852 RNA, 46 ADT, 17,081 ATAC); 6,534 (9,911 RNA, 46 ADT, 16,552 ATAC); and 6,748 (9,859 RNA, 46 ADT, 16,620 ATAC) numbers of cells in each of the four data batches. The cell type information was obtained from the original study and for each of the four data batches the number of cell types are 11, 11, 10, and 10.

#### CITE-seq dataset by Stephenson et al

This CITE-seq dataset measures PBMC from healthy individuals and from COVID-19 patients ^15^. Only the data from healthy individuals were used in this study. The raw matrices of RNA and ADT and the annotation of cells to their respective cell types from the original study were downloaded from the EMBL-EBI ArrayExpress database under the accession number E-MTAB-10026, with 30,313 healthy cells from Cambridge medical centre (batch 1) and 64,262 healthy cells from NCL medical centre (batch 2). RNA and ADT in this dataset were filtered by removing those that expressed in less than 1% of the cells and cell types were filtered by removing those that have less than 10 cells. After filtering, there are 30,313 cells from 17 cell types (10,668 RNA, 192 ADT) in batch 1 and 64,257 cells from 16 cell types (10,618 RNA, 192 ADT) in batch 2 of the dataset for downstream analysis.

#### CITE-seq dataset by Hao et al

The raw RNA and ADT matrices from this CITE-seq dataset generated by Hao et al. from PBMC 2 were downloaded from NCBI GEO under the accession number GSE164378. The dataset contains two batches and the cells in both batches were annotated to 31 cell types. As the above, RNA and ADT in this dataset were filtered by removing those that expressed in less than 1% of the cells and cell types were filtered by removing those that have less than 10 cells. This resulted in 67,090 cells (11,451 RNA, 228 ADT) in batch 1 and 94,674 cells (12,347 RNA, 228 ADT) in batch 2 of the dataset.

#### CITE-seq dataset by Ramaswamy et al

The raw RNA and ADT matrices of PBMC from three healthy donors in this CITE-seq dataset generated by Ramaswamy et al. ^13^ were downloaded from NCBI GEO under the accession number GSE166489. Each patient sample corresponds to one data batch. After filtering RNA and ADT expressed in less than 1% of the cells and discarding cell types that have fewer than 10 cells, we obtained 8,641 cells and 26 cell types in batch 1 (11,062 RNA, 189 ADT), 9,523 cells and 26 cell types in batch 2 (10,801 RNA, 189 ADT), and 10,410 cells and 28 cell types (11,039 RNA, 189 ADT) in batch 3 of this dataset.

#### SHARE-seq dataset

The SHARE-seq data that measures RNA and ATAC from matched cells in mouse skin samples were downloaded from NCBI GEO under the accession number GSE140203 ^16^. The dataset contains raw count of RNA and ATAC of cells annotated to 22 cell types. Similar to the above, we first removed peaks with no expression across cells, and then summarised the ATAC data from peak level into gene activity scores using the ‘CreateGeneActivityMatrix’ function in Seurat. We filtered out RNA and ATAC quantified in fewer than 1% of the cells and cell types that have less than 10 cells, resulting in a dataset with 32,231 cells (8,926 RNA, 14,034 ATAC) for the subsequent analyses.

### Matilda design

#### Multi-task learning architecture

The multi-task neural networks in Matilda consist of multimodality-specific encoders and decoders in a variational autoencoder (VAE) component for data simulation and a fully-connected classification network for cell type classification. The encoders in the VAE component are shareable for both data simulation and classification tasks, and consist of one learnable point-wise parameter layer and one fully-connected layer to the input layer. Because ADT modality has significantly fewer features than RNA and ATAC modalities, we set empirically, based on model selection, the numbers of neurons for encoders of RNA, ADT, and ATAC modalities to be 185, 30, and 185, respectively. To learn a latent space that integrates the information from across modalities, we concatenated the output from the encoder trained from each data modality to perform joint learning using a fully-connected layer with 100 neurons, followed by a VAE reparameterization process ^11^. Next, the fully-connected layer of the latent space is split into two branches with one branch fed into the decoders and the other branch fed into the fully-connected classification network. For the decoder branch, it consists of multiple decoders each corresponds to an input data modality. Each decoder consists of one fully-connected layer to the output layer that has the same number of neurons as the features in the corresponding data modality. For each fully-connected layer in the VAE component, batch normalisation ^25^, shortcut ^26^ were utilised in the model. ReLU activation was used in all fully-connected layers except in the reparameterization process. Dropout (r=0.2) was utilised only for fully-connected layers in encoders. For the classification branch, it consists of the latent space as input to a fully-connected layer with a dimension equal to the number of cell types in the training data. The fully-connected layer outputs a probability vector for cell type prediction through a SoftMax function.

#### Loss function

Let *X* be the single-cell multimodal omic data from *N* modalities, the VAE component of Matilda contains two procedures: (1) the encoders encode each modality in the data *X* individually, and concatenate them for joint learning. This process projected the high-dimensional *X* into a lowdimensional latent space. We denote the posterior distribution of this process as *q_θ_*(*z|X*), where *θ* is the learnable parameter of the neural network in this procedure. (2) the decoders reconstruct the lowdimensional latent space to the high-dimensional original data space. We denote the posterior distribution of this process as *p_φ_*(*X|z*), where *φ* is the learnable parameter of the neural network in this procedure. The loss function of the data simulation component can be represented as the negative loglikelihood with a regularizer:

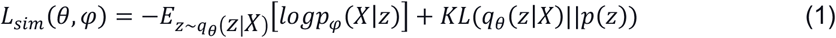

The first term is the reconstruction loss using the expectation of negative log-likelihood. This term encourages the decoder to learn to reconstruct the original data *X* using the low-dimensional representation *z*. The second term is the Kullback-Leibler (*KL*) divergence between the encoder’s distribution *q_θ_*(*z|X*) and *p*(*z*), where *p*(*z*) is specified as a standard Normal distribution as *p*(*z*)~*N*(0,1). This divergence measures the information loss when using *q_θ_*(*z|X*) to represent *p*(*z*). The encoder network parameters are in turn optimised using stochastic gradient descent via back-propagation, which is made possible by the reparameterization trick ^11^.

For the loss function of the classification component, we use cross-entropy loss with label smoothing ^27^. Label smoothing is a regularizer technique, which replaces one-hot real label vector *y_real_* with a mixture of *y_real_* and the uniform distribution:

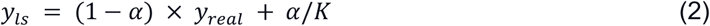

where *K* is the number of label classes, and *α* is a hyperparameter that determines the amount of smoothing. Then, the classification loss can be represented as:

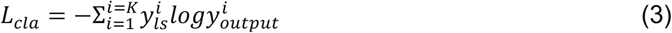

where 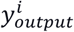 is the predicted label for the *i^th^* cell.

To learn Matilda, we combined the simulation loss and classification loss to give the following overall loss function:

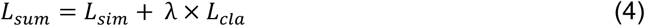

where *λ* is a weighting coefficient that determines the importance of the classification term against the data simulation term from Matilda.

#### Data augmentation and balancing strategy

During the model training process, Matilda performs data augmentation and balancing using simulated data from the VAE component. Specifically, Matilda first ranks the cell types in the training dataset by the number of cells in each type. The cell type corresponding to the median number is used as the reference and those that have smaller numbers of cells are augmented to have the same number of cells as the median using VAE simulated single-cell multimodal data for each cell type. Cell types that have larger numbers of cells than the median number are randomly down-sampled to match the median number of cells as well. This strategy helps Matilda to mitigate imbalanced cell type distribution in the data ^28^ and better learn the molecular features of under-represented and rare cell types.

#### Joint feature selection from multiple modalities

Leveraging its neural network architecture, Matilda implements a novel approach to detect the most informative features simultaneously from each of all data modalities. Specifically, to assess the importance of each feature, the trained model was used for back-propagation of the partial derivatives from the output units of the classification network to the input units of the encoders, where each input unit represents an individual feature from a given modality in the input data *X*. The importance score of each input feature of each cell is determined by approximating the integral gradients of the model’s output to its input:

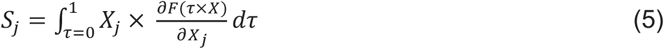

where *F* represents the classification branch of the multi-task neural networks, and 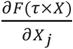 is the gradient of *F*(*X*) along with the *j^th^* feature. We aggregated these derivatives across cells within each cell type. These aggregated gradients indicate the importance of each feature from each data modality in predicting each cell type. The top-ranked features from each cell type can be selected based on their aggregated derivatives for subsequent analyses.

#### Matilda model training

Matilda adopts a two-step training strategy. In the first step, i.e., before augmentation and balancing, we train a network from scratch. In the second step, i.e., after augmentation and balancing, we inherit the weights from the first step as the initial value and fine-tune the networks using augmented and balanced data. Several key hyper-parameters may impact the performance of Matilda. These include the number of layers in the neural networks, the number of neurons in each layer, the parameter *λ* that balances the VAE data reconstruction and cell type classification in the multi-tasking learning, and other parameters such as learning rate, number of epochs, batch size, and dropout rate. To optimise these hyper-parameters, we used the training datasets of CITE-seq, SHARE-seq, and TEA-seq to evaluate the model performance with different parameter combinations based on measurements including (a) the distance between the umap of simulated data and real data and (b) the classification accuracy before and after data augmentation. These allowed us to determine the following Matilda settings that were used in subsequent experiments. Specifically, for both steps in the training process, batch size was set to 64 cells in learning from all datasets. The epoch was set to 30 for all datasets except the CITE-seq dataset generated by Hao et al. (GSE164378) which contains the largest number of cells. Since large datasets do not need many training epochs for the neural networks to converge, we set this to 10 for this CITE-seq dataset (GSE164378) for improving training efficiency. The parameter *λ* for balancing loss function in multitasking learning was empirically set to 0.1 for all datasets and the parameter *α*. in label smoothing was set to 0.1 according to ^29^. In the first stage, we empirically determined the learning rate of 0.02 in the training process. In the second stage, we fine-tuned the networks with an initial learning rate of 0.02 for the first half of epochs and 0.002 for the second half of epochs. In Matilda, all input data modalities were normalised by the ‘NormalizeData’ function in Seurat ^14^ and then scaled using a z-score transformation to a similar range.

### Settings for other classification methods

#### CHETAH

Raw count matrices of RNA modality from each dataset were used as input for CHETAH (v1.8.0) ^6^ and the function ‘CHETAHclassifier’ was used to perform cell type classification, following the author’s tutorial (https://github.com/jdekanter/CHETAH).

#### scmapCell

Raw count matrices of RNA modality from each dataset were first normalised using ‘NormalizeData’ function in Seurat and then used as input for scmap (v1.14.0) ^30^ as suggested (https://github.com/hemberg-lab/scmap). By default, the top 500 most informative genes were used and the function ‘scmapCell2Cluster’ annotates cells in the query dataset to their respective cell types based on the reference data.

#### scClassify

Raw count matrices of RNA modality from each dataset were first normalised using the ‘NormalizeData’ function in Seurat and then used as input for scClassify (v1.4.0) ^5^. The default parameters, e.g., tree=“HOPACH”, algorithm=“WKNN”, selectFeatures=“limma”, similarity=“pearson”, were used as suggested in the pipeline (https://github.com/SydneyBioX/scClassify).

#### singleCellNet

Raw count matrices of RNA modality from each dataset were first normalised using the ‘NormalizeData’ function in Seurat and then used as input for singleCellNet (v0.1.0) ^31^. ‘scn_train’ function with the default parameters of nTopGenes=10, nRand=70, nTrees=1000, nTopGenePairs=25, dLevel=“newAnn”, colName_samp=“cell” was used for training the model. The trained models were subsequently used for predicting the cell types for cells in the query data using ‘scn_predict’ and ‘assess_comm’ with default parameters (https://github.com/pcahan1/singleCellNet).

#### CelliD

The raw count matrices of RNA modality from each dataset were used as input for CelliD (v1.0.0) ^32^. Following the author’s pipeline (https://github.com/RausellLab/CelliD), we use the function ‘RunMCA’ to perform Multiple Correspondence Analysis (MCA) dimension reduction for both reference and query data. Then extract gene signatures in each cell type using the function ‘GetGroupGeneSet’ with default parameters dims = 1:50, n.features = 200, group.by=“cell.type”. The cell-to-cell matching and label transferring across data were generated using the function ‘RunCellHGT’.

#### scID

Raw count matrices of RNA modality from each dataset were first normalised using the ‘NormalizeData’ function in Seurat and then used as input for the R package scID (v2.2) ^33^. Following the author’s tutorial (https://github.com/BatadaLab/scID), we used the function ‘scid_multiclass’ with default parameters for identifying cell types in the query datasets.

### Settings for other simulation methods

#### SPARSim

Following the author’s pipeline (https://gitlab.com/sysbiobig/sparsim), raw count matrices of RNA modality from each dataset were used as input for SPARSim (v0.9.5) ^17^. Data were first normalised using the ‘scran_normalization’ function in SPARSim package and data parameters were estimated by ‘SPARSim_estimate_parameter_from_data’ function. The function ‘SPARSim_simulation’ was then used for generating simulated data.

#### cscGAN

Following the author’s pipeline (https://github.com/imsb-uke/scGAN), raw count matrices of RNA modality from each dataset were first normalised using the ‘process_files’ function in cscGAN (Github version 988ad95) ^18^. Default parameters and training iteration of 6000 was used for model training and the ‘run_exp’ function was used for generating simulated data from the trained model.

### Settings for other dimension reduction methods

#### Seurat

Seurat package (v4.1.0) ^14^ was used for dimension reduction of all CITE-seq datasets. The raw count matrices of RNA and ADT were used as input, which were then normalised by the ‘NormalizeData’ function in Seurat. By default, the top 2,000 most variable genes were selected from RNA modality by ‘FindVariableFeatures’ function and data are subsequently scaled by ‘ScaleData’ function. Data from ADT modality were processed similarly as those of RNA modality, except using parameters of normalization.method=“CLR” and margin=2 in the in ‘NormalizeData’ function, as suggested in the author’s pipeline (https://satijalab.org/seurat/). PCA was performed using the ‘runPCA’ function and the function ‘FindMultiModalNeighbors’ integrates RNA and ADT modalities using the PCA results. The joint visualisation of RNA and ADT were generated using ‘wnn.umap’ function.

#### totalVI

The totalVI procedure implemented in the scvi-tools package (v0.15.0) ^4^ was used for dimension reduction of all CITE-seq datasets. Following the author’s tutorial (https://github.com/scverse/scvi-tools), the raw count matrices of RNA and ADT were first normalised using the ‘normalize_total’ and ‘log1p’ functions and then the top 4,000 most variable genes were selected using the ‘highly_variable_genes’ function. The data were subsequently used as input for model training using ‘scvi.model.TOTALVI.setup_anndata’, ‘scvi.model.TOTALVI’, and ‘train’ functions in scvi-tools. The latent space of RNA and ADT modalities was generated using the ‘get_latent_representation’ function.

#### Conos

Conos package (v1.4.5) ^4^ was used for dimension reduction of the SHARE-seq dataset. Following the author’s pipeline (https://github.com/kharchenkolab/conos), the raw count matrices of RNA and ATAC were normalised by the ‘basicP2proc’ function in pagoda2 package (v1.0.8), where recommended parameters of n.odgenes=3e3, nPcs=30, min.cells.per.gene=-1, make.geneknn=FALSE, and n.cores=1 were used. Next, the joint graph was built using buildGraph with k=15, k.self=5, k.self.weigh=0.01, ncomps=30, n.odgenes=5e3, and space=“PCA” in Conos. The joint visualisation of RNA and ATAC were generated using ‘largeVis’ in function ‘embedGraph’ with default parameter alpha=1/2.

#### MultiVI

The MultiVI procedure implemented in the scvi-tools (v0.15.0) ^20^ was used for dimension reduction of the SHARE-seq dataset. Following the author’s pipeline (https://github.com/scverse/scvi-tools), the raw count matrices of RNA and gene activity score matrices from ATAC and the paired matrix of RNA and ATAC were used as input. These data were first concatenated using the ‘organize_multiome_anndatas’ function in scvi-tools and then used for model training using ‘scvi.model.MULTIVI.setup_anndata’, ‘scvi.model.MULTIVI’ and ‘train’ functions in scvi-tools. The latent space of RNA and ATAC modalities was generated using the ‘get_latent_representation’ function.

### Settings for other feature selection methods

We performed feature selection from the RNA modality of each dataset using a collection methods: (i) simple one-sided *t*-test and Wilcoxon rank sum test, (ii) popular methods based on differential expression analysis including Limma (v3.48.3) ^7^ and MAST (v.1.2.1) ^8^, and (iii) methods based on maximising classification performance including logistic regression (LR) and receiver operating curve (ROC) implemented in the ‘FindMarkers’ function in Seurat 2.

### Performance evaluation

#### Cell type classification evaluation

We evaluated the accuracy of a cell type classification model by calculating their average accuracy as the sum of the accuracy in all cell types divided by the number of cell types in a dataset. The average accuracy of all cell types accounts for the performance of a classification model in both the major and minor cell types. We used two pipelines, referred to as “intradataset” and “inter-dataset” classification, for cell type classification model evaluation. While intradataset classification splits training and test data from one batch of a dataset, inter-dataset classification splits training and test data from different batches in a dataset. For intra-dataset classification, we performed five-fold cross-validation repeated five times with different seeding on each batch of each dataset. For inter-dataset classification, we select the common features and cell types from different batches in the same dataset with different data batches and train on one batch and test on another batch.

#### Simulation evaluation

We used the correlation heatmaps to visualise the correlation structure of select features in each data modality of each dataset. Specifically, we first applied the functions ‘modelGeneVar’ and ‘getTopHVGs’ from the scran R package (v1.20.1) ^34^ to select the top 100 high variable genes (HVGs) based on their variability calculated from each data modality in each dataset (except the ADT modality of the TEA-seq dataset) and then calculated pairwise Pearson’s correlation coefficients from these HVGs across all cells in each dataset. Since the ADT modality of the TEA-seq dataset only contains 46 ADTs, we used all of them in the correlation analysis and heatmap visualisation. For comparison to other simulation methods in RNA modality, we used the same visualisation methods as above for each simulation method and also quantified the performance of each simulation method by calculating the overall Pearsons’s correlation of real and simulated data and represented these as boxplots.

#### Dimension reduction evaluation

We used the performance of a simple *k*-means clustering algorithm to assess cell type clustering on dimension reduced dataset generated from each modality integration and dimension reduction method. Similar to cell type classification, we used intra-dataset and inter-dataset for assessing cell type clustering. In particular, we used the latent space of the test dataset obtained either from five-fold cross-validation or a data batch for cell type clustering and compared the concordance between the clustering output and the cell type labels from their original studies. The fivefold cross-validation procedure in the intra-dataset clustering was repeated five times with different seeding. We assessed the clustering concordance using four evaluation metrics, including Adjusted Rand Index (ARI), Normalized Mutual Information (NMI), Fowlkes-Mallows index (FM), and Jaccard index (Jaccard). Briefly, let *N* be the number of cells in the dataset, *u* = {*U*_1_, *U*_2_, …, *U_R_*} be the cell type annotation from the original study, and *v* = {*V*_1_, *V*_2_, …, *V_C_*} be the partition generated by clustering, the pairs between *u* and *v* can be classified into one of the four types: (i) *N*_11_: the number of pairs that are in the same partition in both *u* and *v;* (ii) *N*_00_: the number of pairs that are in different partitions in *u* and *v;* (iii) *N*_01_: the number of pairs that are in the same partition in *u* but in different partitions in *v;* (iv) *N*_10_: the number of pairs that are in different partitions in *u* but in the same partition in *v*. Given the above notation, we defined the ARI, NMI, FM, Jaccard metrics as follows:

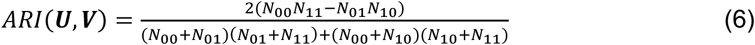

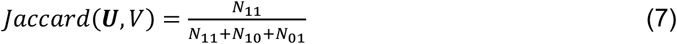

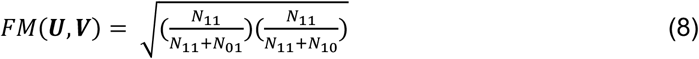

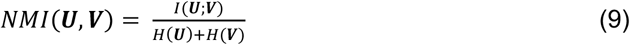

where *I*(***U**; **V***) is the mutual information between ***U*** and ***V***, defined as

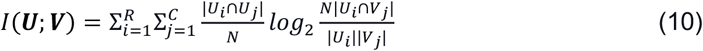

and *H*(·) is the entropy of partitions, in which *H*(***U***) and *H*(***V***) are calculated

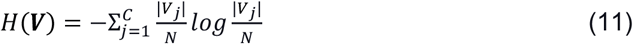

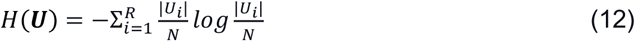

#### Feature selection evaluation

We used the classification of each cell type to evaluate the performance of features selected for that cell type. Specifically, we used a ‘one-vs-all’ procedure in that we classified each cell type against all remaining cell types using the top 100 features selected for that cell type from different feature selection methods. Note that only Matilda selected features from all data modalities whereas the other feature selection methods were designed for analysing gene expression data and thus used only to select features from RNA modality of each dataset. The classification accuracy for each cell type was calculated using the ‘intra-dataset’ procedure in that feature selection was conducted on training datasets and their utility/effectiveness in cell type classification were verified on test datasets generated from five-fold cross-validation repeated five times.

#### Running time evaluation

We evaluated running time on a server with AMD(R) Ryzen processor CPU (16 cores and 64 Gb total memory) and one RTX3090 graphics processing unit. We used the CITE-seq datasets generated by Hao et al. (GSE164378) and Ramaswamy et al. (GSE166489) to benchmark the running time, given the large numbers of cells in these two datasets. To evaluate the impact of the number of cells from the training datasets, we kept the number of cells to 2k in the test dataset and varied the number of cells in the training datasets from 1k, 2k, 3k, 5k, 10k, 20k, to 30k. Similarly, to evaluate the impact of the number of cells from test dataset, we kept the number of cells in the training dataset to 2k and varied the those in the test datasets from 1k, 2k, 3k, 5k, 10k, 20k, to 30k as above. The elapsed run time was evaluated by the R function ‘system.time()’ and Python function ‘time.time()’ for methods implemented using R and Python, respectively.

## Acknowledgements

We think the intellectual feedback from the Computational Systems Biology Group at Children’s Medical Research Institute (CMRI) and engagement from the colleagues at the School of Mathematics and Statistics, The University of Sydney and Sydney Precision Bioinformatics Alliance. This work is funded by National Health and Medical Research Council (NHMRC) Investigator Grant (1173469) to P.Y.

## Author contributions

Conceptualization, C.L., P.Y.; Methodology, C.L., P.Y.; Investigation, C.L., H.H., P.Y.; Data generation, C.L., H.H.; Writing – Original Draft, C.L., P.Y.; Writing – Review & Editing, all authors; Supervision, P.Y.; Funding Acquisition, P.Y.

## Data availability

All the datasets used in this study are publicly available. The “TEA-seq dataset” was downloaded from NCBI GEO under the accession number GSE158013. The “CITE-seq dataset by Stephenson et al” was downloaded from the EMBL-EBI Array Express database under the accession number E-MTAB-10026. The “CITE-seq dataset by Hao et al” was downloaded from NCBI GEO under the accession number GSE164378. The “CITE-seq dataset by Ramaswamy et al” was downloaded from NCBI GEO under the accession number GSE166489. The “SHARE-seq” was downloaded from NCBI GEO under the accession number GSE140203.

## Code availability

Matilda was implemented using PyTorch (version 1.9.1) with code available at https://github.com/liuchunlei0430/Matilda.

