## supplementary information for "Multi-task learning from single-cell multimodal omics with Matilda"

### Supplementary Figures

**a**

| Dataset | Protocol | Modalities | Species | Tissue | Batch | # cell type | # cell |
|---|---|---|---|---|---|---|---|
| Swanson et al. (GSE158013) | TEA-seq | RNA, ADT, ATAC | Human | PBMC | B1 | 11 | 6,310 |
| | | | | | B2 | 11 | 6,545 |
| | | | | | B3 | 10 | 6,534 |
| | | | | | B4 | 10 | 6,748 |
| Ramaswamy et al. (GSE166489) | CITE-seq | RNA, ADT | Human | PBMC | B1 | 26 | 8,641 |
| | | | | | B2 | 26 | 9,523 |
| | | | | | B3 | 28 | 10,410 |
| Hao et al. (GSE164378) | CITE-seq | RNA, ADT | Human | PBMC | B1 | 31 | 67,090 |
| | | | | | B2 | 31 | 94,674 |
| Stephenson et al. (E-MTAB-10026) | CITE-seq | RNA, ADT | Human | PBMC | B1 | 17 | 30,313 |
| | | | | | B2 | 16 | 64,257 |
| Ma et al. (GSE140203) | SHARE-seq | RNA, ATAC | Mouse | Skin | NA | 22 | 32,231 |

**b**

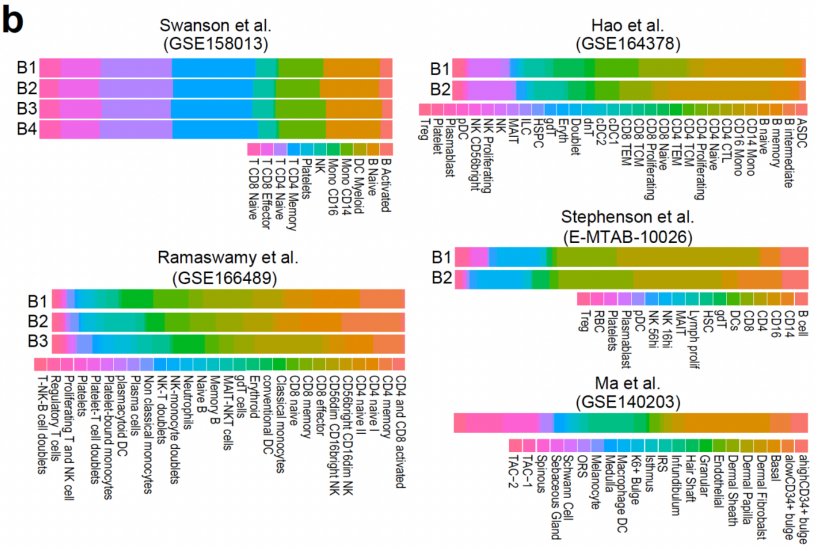


**Supplementary Figure 1**: Single-cell multimodal omics datasets included in this study. (a) A summary of each single-cell multimodal omics dataset including the experimental protocol used, modalities profiled, biological sample information, and information of batch, number of cell types, and number of cells for each dataset. (b) Cell type annotation and their proportions in each dataset and batch.

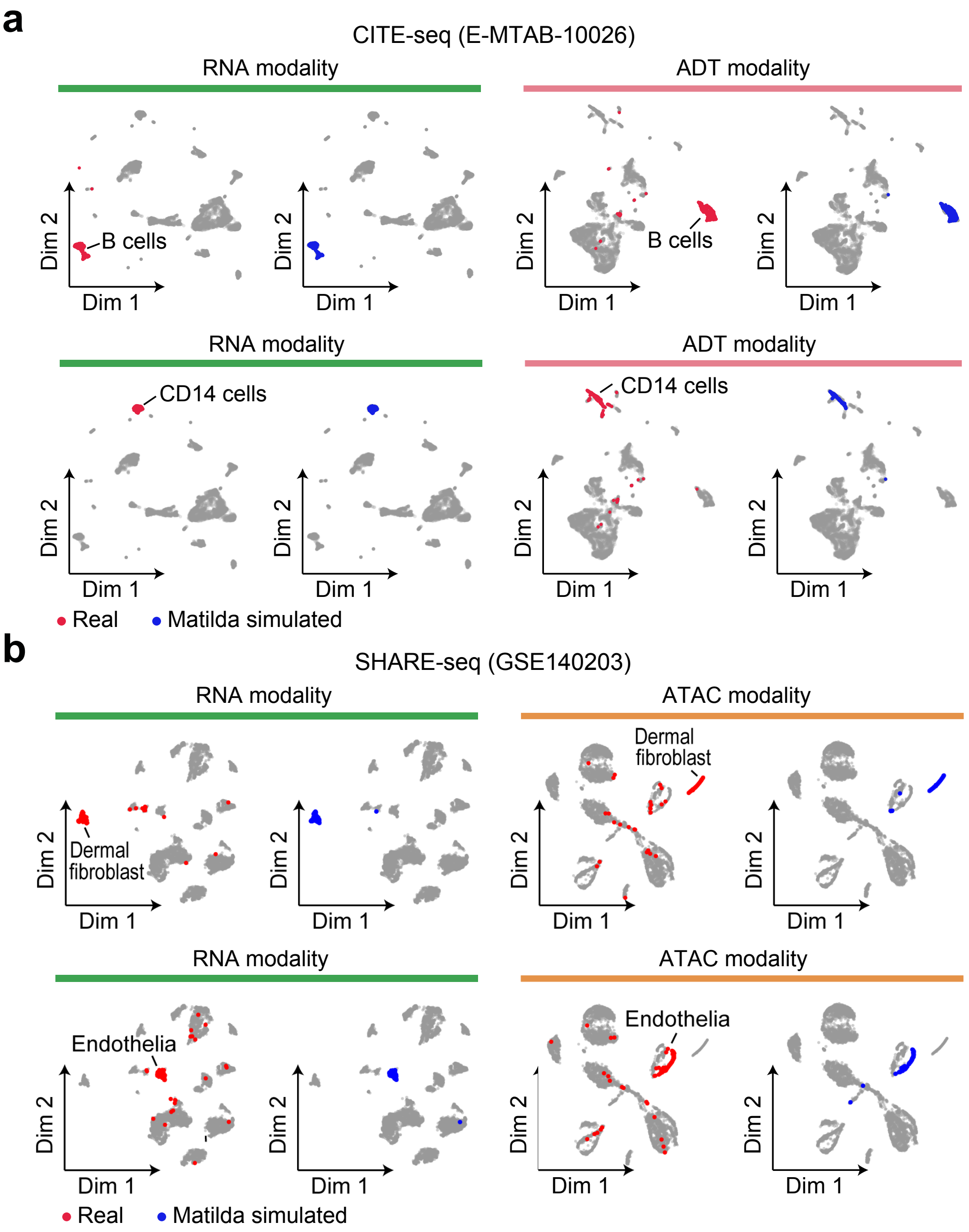


**Supplementary Figure 2**: UMAP visualization of cell-type-specific simulations of single-cell multimodal data using Matilda. (a) Real (red) and Matilda simulated (blue) B cells (upper panels) and CD14 cells (lower panels) visualized on UMAPs using RNA modality (left panels) and ADT modality (right panels). (b) Real (red) and Matilda simulated (blue) dermal fibroblasts (upper panels) and endothelial cells (lower panels) visualized on UMAPs using RNA modality (left panels) and ATAC modality (right panels).

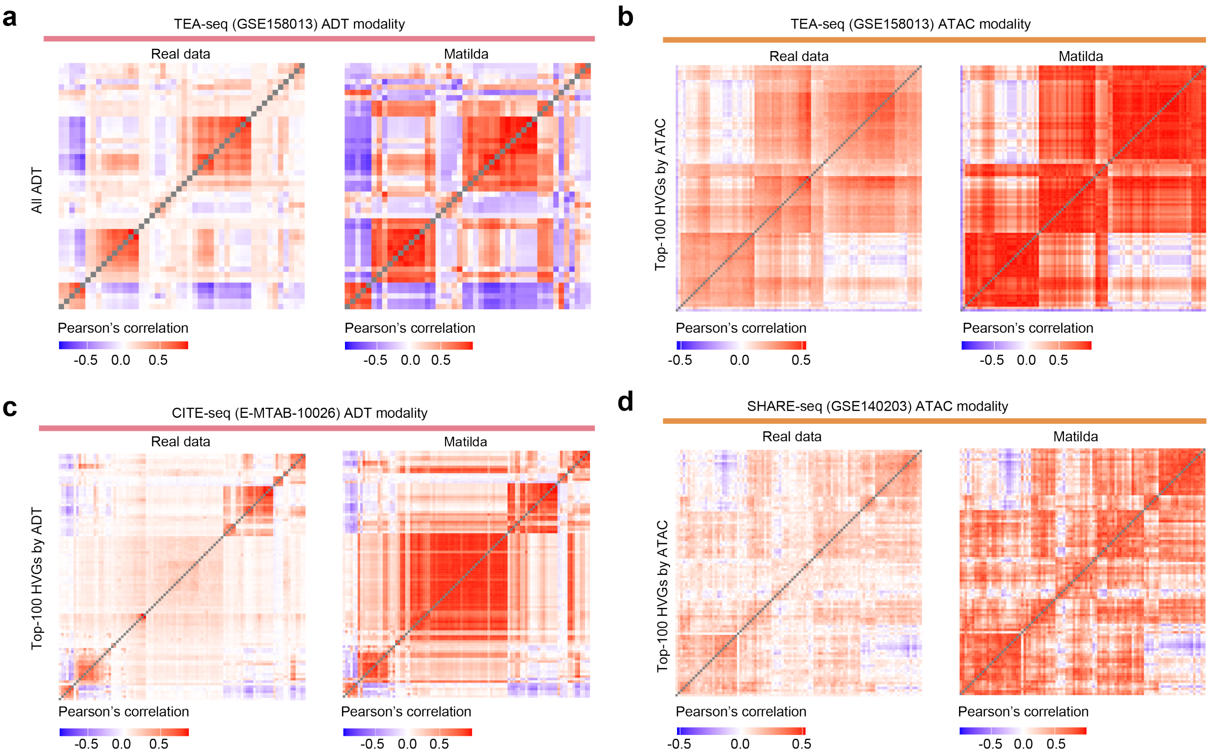


**Supplementary Figure 3**: Heatmap visualization of the correlation structure of real and Matilda simulated data. (a) Correlation structures of all ADTs in real (left) and Matilda simulated (right) TEA-seq (GSE158013) data using ADT modality. (b) Correlation structure of the top 100 HVGs selected by ATAC modality in real (left) and Matilda simulated (right) TEA-seq (GSE158013) data using ATAC modality. (c) Correlation structures of the top 100 HVGs selected by ADT modality in real (left) and Matilda simulated (right) CITE-seq (E-MTAB-10026) data using ADT modality. (d) Correlation structure of the top 100 HVGs selected by ATAC modality in real (left) and Matilda simulated (right) SHARE-seq (GSE140203) data using ATAC modality.

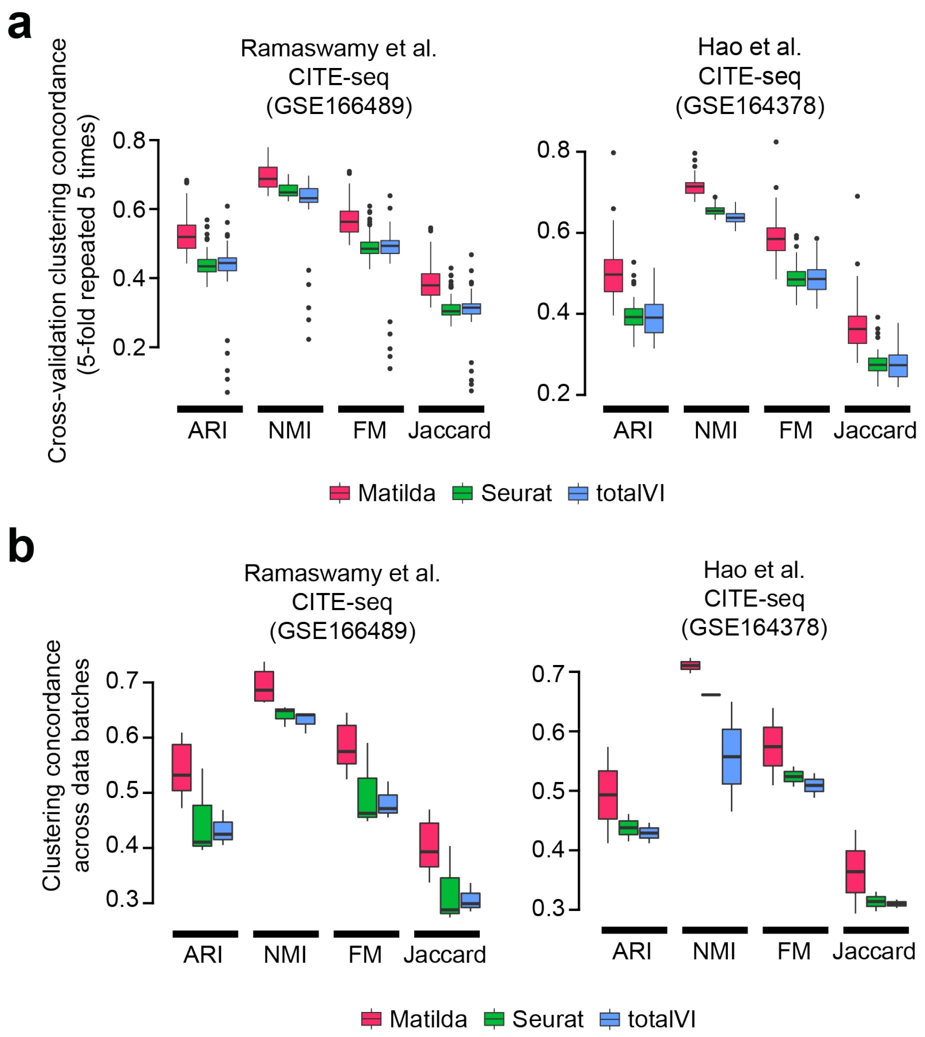


**Supplementary Figure 4**: Quantification of joint multimodal dimension reduction results from CITE-seq data using Matilda, Seurat, and totalVI. (a) Quantification of k-means clustering concordance using dimension reduced data from Matilda, Conos, and MultiVI with cell-type annotation from the original study by ARI, NMI, FM, and Jaccard index. 5-fold cross-validation repeated five times with different random seedings were used for the comparison. (b) Same as (a) but comparing the performance across data batches. Centre line, median; box limits, upper and lower quartiles; whiskers, 1.5x interquartile range; points, outliers.

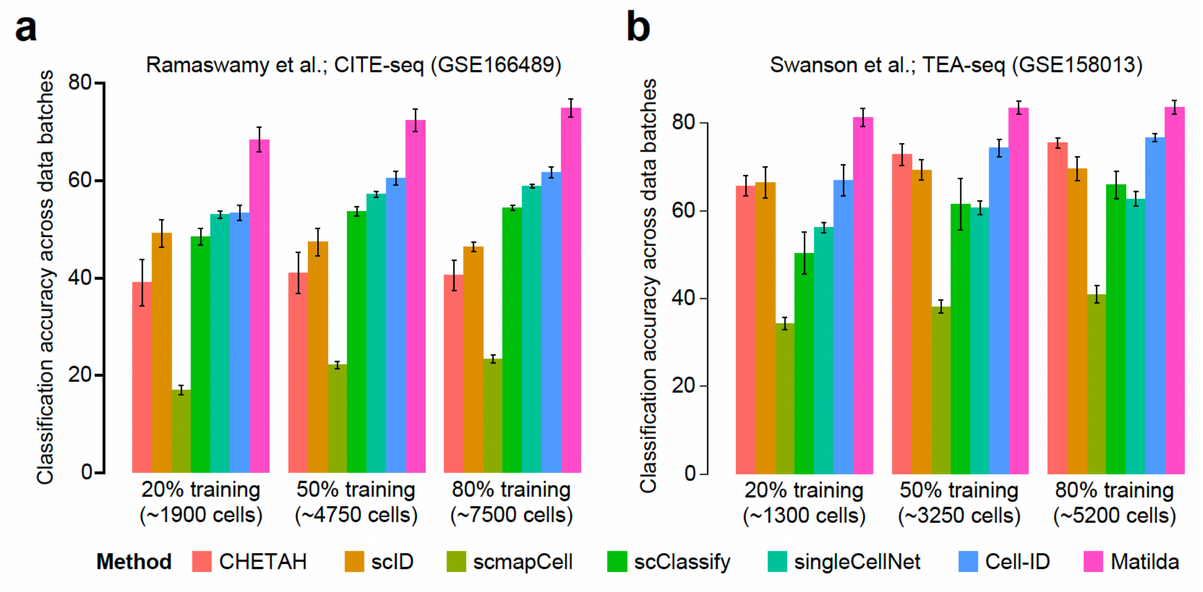


**Supplementary Figure 5**: Benchmarking performance of cell type classification. (a) Cell type classification of each method using CITE-seq (GSE166489) dataset. (b) Cell type classification of each method using TEA-seq (GSE158013) dataset. Classification models from each method are trained on different numbers of cells subsampled from the original data to investigate the impact of training data size on classification performance. Error bar, SD.

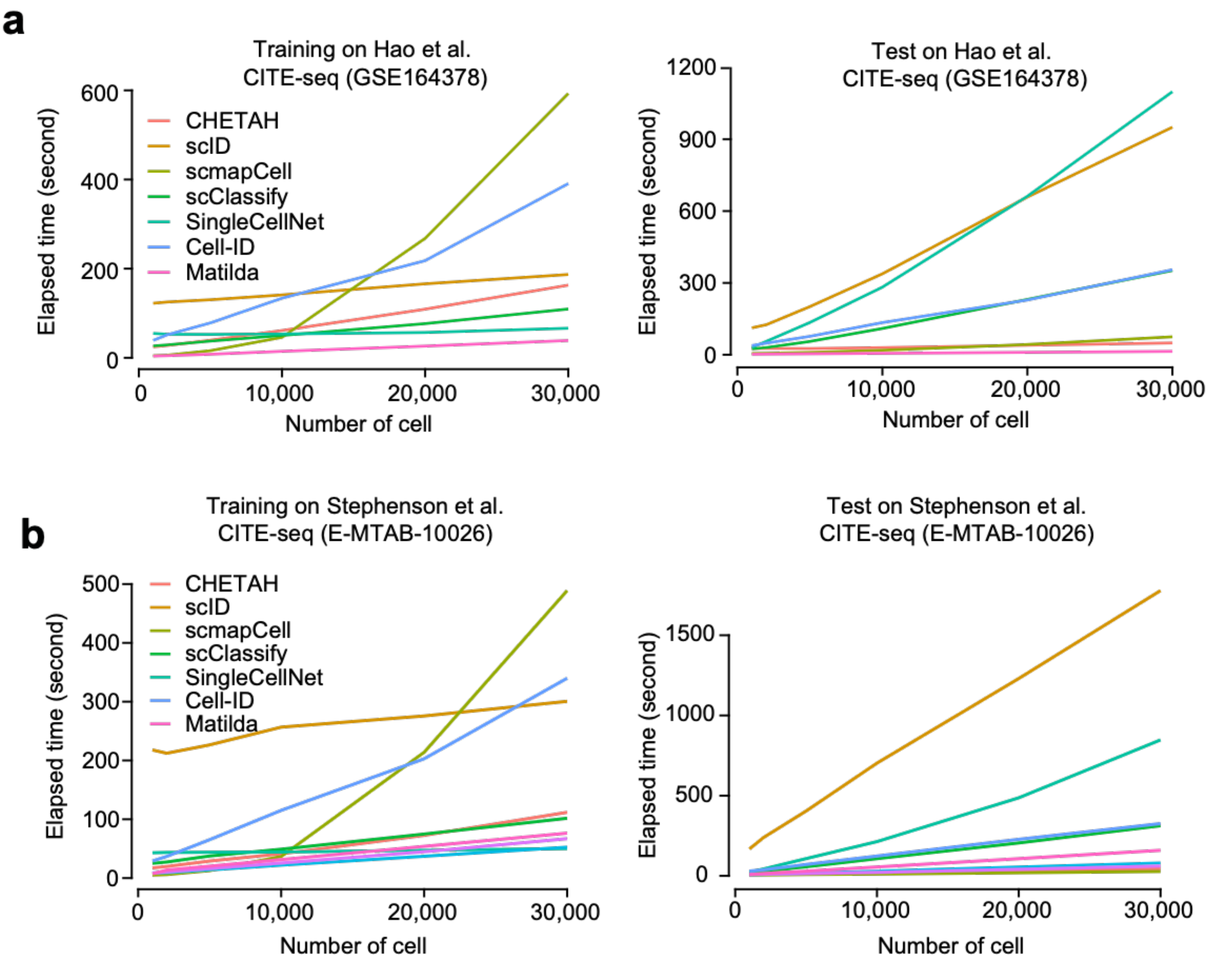


**Supplementary Figure 6**: Model training and classification time on data with different number of cells for each method using (a) GSE164378 and (b) E-MTAB-10026 datasets.

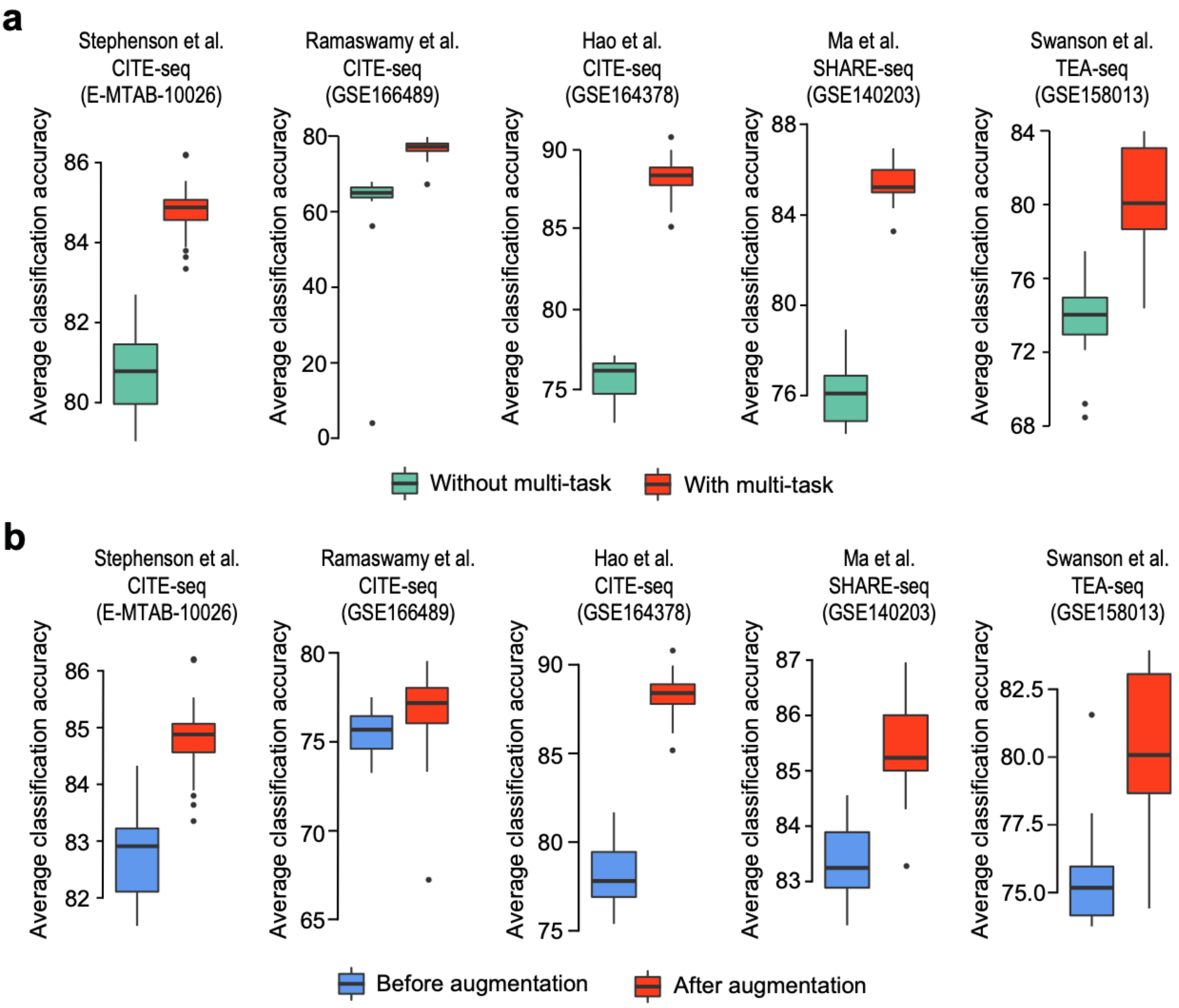


**Supplementary Figure 7**: Evaluation of multi-task learning and data augmentation and balancing in Matilda. (a) Cell type classification accuracy comparison of Matilda with and without multi-task learning in which models for data simulation and cell type classification are trained separately. (b) Cell type classification accuracy comparison of Matilda before and after training the model with data augmentation and balancing. Centre line, median; box limits, upper and lower quartiles; whiskers, 1.5x interquartile range; points, outliers.

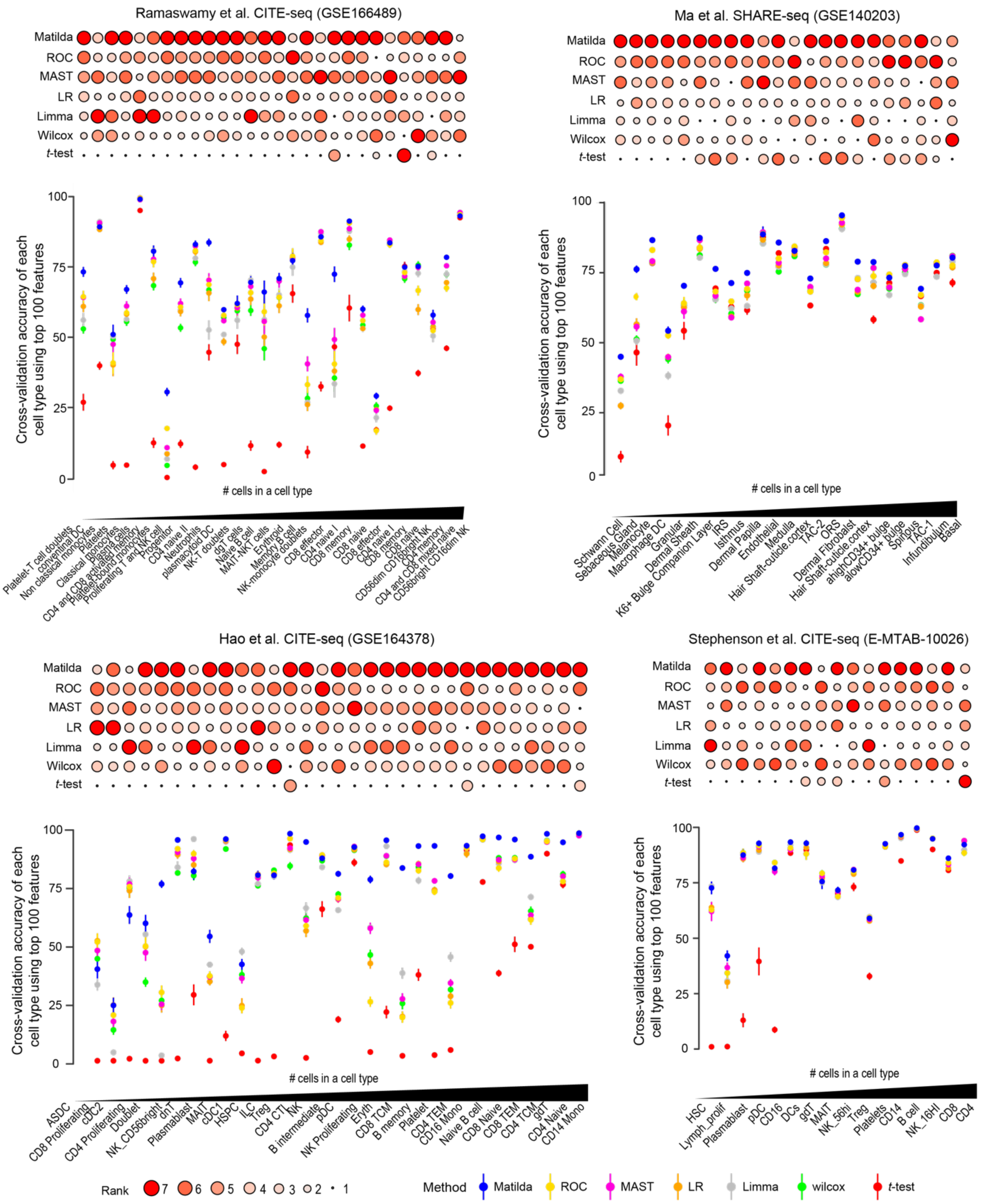


**Supplementary Figure 8**: Classification of each cell type using features selected by different methods for that cell type for GSE166489, GSE140203, GSE164378, and E-MTAB-10026 datasets. Cell types are arranged from low to high based on the number of cells in each cell type. Feature selection methods are also ranked based on the performance of their selected features in classifying each cell type (upper panel). Error bar, SE.
